# Dissection of PRC1 and PRC2 recruitment in Arabidopsis connects EAR repressome to PRC2 anchoring

**DOI:** 10.1101/2020.08.28.271999

**Authors:** Fernando Baile, Wiam Merini, Inés Hidalgo, Myriam Calonje

## Abstract

PcG complexes ensure that every cell in an organism expresses the genes needed at a particular stage, time or condition. However, it is still not fully understood how PRC1 and PRC2 are recruited to target genes in plants. Recent results in Arabidopsis support that PRC2 recruitment is mediated by different TFs. However, it is unclear how all these TFs interact with PRC2 and whether they can also recruit PRC1 activity. Here, by using a system to *in vivo* bind selected factors to a synthetic promoter lacking the complexity of PcG target promoters, we show that while VAL1 binding recapitulates PRC1 and PRC2 marking, the binding of other TFs only render PRC2 marking. Interestingly, all these TFs contain an EAR domain that acts as docking point for PRC2 and HDACs, connecting two different repressive mechanisms. Furthermore, we show that different TFs act synergistically in PRC2 anchoring to maintain a long-term repression.

## Introduction

The evolutionary conserved Polycomb group (PcG) factors are required to maintain gene repression (Ringrose and Paro, 2004; Merini and Calonje, 2015). These factors form multiprotein complexes with different histone modifying activities, including PcG repressive complex 1 (PRC1), which has H2A E3 ubiquitin ligase activity towards lysine 119, 120 or 121 in Drosophila, mammals or Arabidopsis, respectively (Wang et al., 2004; Cao et al., 2005; Bratzel et al., 2010; Yang et al., 2013), and PRC2, which has H3 lysine 27 (H3K27) trimethyltransferase activity (Müller et al., 2002; Makarevich et al., 2006; Mozgova and Hennig, 2015). Despite the conserved activity of these complexes, several data indicate that distinct rules operate for PcG recruitment in the different organisms (Müller and Kassis, 2006; Mendenhall et al., 2010; Xiao et al., 2017).

In Arabidopsis, PRC2 core subunits are well conserved to their animal counterpart (Mozgova et al., 2015); however, PRC1 composition is less conserved (Merini and Calonje, 2015). Although a H2A E3 ubiquitin ligase module containing one AtBMI1 (A, B or C) and one AtRING1 (A or B) protein has been identified (Bratzel et al., 2010; Sanchez-Pulido et al., 2008), homologs for other PRC1 components are missing and instead several plant-specific proteins seem to play PcG functions (Merini and Calonje, 2015; Calonje, 2014). Distribution analysis of H2AK121ub and H3K27me3 peaks in Arabidopsis showed that both marks are generally targeted to gene regions, although H3K27me3 peaks are longer than H2AK121ub peaks. In addition, this analysis revealed that despite H2AK121ub marks frequently co-localize with H3K27me3, there are also genes only marked with H3K27me3 or H2AK121ub (Zhou et al., 2017).

Concerning PcG recruitment to target genes in Arabidopsis, a high number of transcription factors (TFs) has been related to PRC2 tethering. Among these factors are the *GAGA* motif binding proteins BASIC PENTACYSTEINE (BPC) 1-6 (Hecker et al., 2015a; Xiao et al., 2017), the *TELOBOX* motif binding proteins ARABIDOPSIS ZINC FINGER 1 (AZF1), ZINC FINGER OF ARABIDOPSIS THALIANA 6 (ZAT6) (Xiao et al., 2017) and TELOMERE-REPEAT-BINDING FACTOR (TRB)1/2/3 (Zhou et al., 2018), the MYB TF ASYMMETRIC LEAVES 1 (AS1) (Lodha et al., 2013), the C2H2 TFs SUPERMAN (SUP) (Xu et al., 2018) and KNUCKLES (KNU) (Sun et al., 2019), and the MADS-box TFs FLOWERING LOCUS C (FLC) and SHORT VEGETATIVE PHASE (SVP) (Wang et al., 2014; Richter et al., 2019). Furthermore, it has been recently shown that certain genomic fragments located at several PcG targets, which contain binding sites for a wide variety of TF families, can recruit PRC2, thus, functioning as Drosophila Polycomb Recruiting Elements (PREs) (Xiao et al., 2017). In addition, localization analyses of H2AK121ub and H3K27me3 marks in WT and PcG mutants showed that levels of H3K27me3 are substantially reduced in the PRC1 mutant *atbmi1abc* (Zhou et al., 2017), indicating that PRC1 also plays a role in PRC2 recruitment.

Unlike PRC2, the recruitment of PRC1 H2A E3 ubiquitin ligase module has so far only been associated to VIVIPAROUS1/ABI3-LIKE (VAL)1/2 proteins (Yang et al., 2013; Qüesta et al., 2016), which is surprising given the number of TFs involved in PRC2 recruitment and the apparent dependence of PRC1 for H3K27me3 marking. Thus, despite recent advances in understanding PcG recruitment in plants, there are still many unknowns. For instance, it is still far from clear whether the recruitment of one complex is required for the recruitment of the other, how PRC2 can interacts with such a diversity of TFs, and whether these interactions take place independently or in parallel. In addition, as there are genes marked with H2AK121ub/H3K27me3, H2AK121ub or H3K27me3 (Zhou et al., 2017), it is unknown whether this differential marking depends on different recruiting factors and, in that case, if these factors can function synergistically at some target genes.

To address all these questions, we developed a system to *in vivo* mediate the binding of selected factors to a synthetic promoter lacking the *cis* regulatory elements involved in PcG recruitment, allowing us to investigate their role under controlled conditions. Our results show that VAL1 can recapitulate PRC1 and PRC2 marking. However, while PRC1 recruitment is directly mediated by interaction with VAL1, PRC2 tethering involves both VAL1 and PRC1. Interestingly, we also found that PRC2 can be recruited independently of PRC1 by TFs from different families that contain an EAR domain as a common feature. We show that the EAR domain, through its interaction with TOPLESS (TPL)/TPL-RELATED (TPR)1-4 corepressors or the SIN3-associated protein 18 (SAP18), acts as a docking point for both PRC2 and HISTONE DEACETHYLASE COMPLEXES (HDACs). Furthermore, we found that different TFs could act synergistically in PRC2 recruitment, leading to increased levels of H3K27me3 at target genes. Our results not only unveil how the different PcG complexes are recruited to target genes, but also how different histone modifying activities are coupled to promote gene repression in Arabidopsis.

## Results

### VAL1 acts as a platform for simultaneous recruitment of PRC1, PRC2 and HDACs

VAL1/2 TFs have been involved in both PRC1- and PRC2-mediated repression (Yang et al., 2013; Qüesta et al., 2016; Chen et al., 2018; Jing et al., 2019; Zeng et al., 2020). VAL factors contain a B3 DNA-binding domain that specifically recognizes RY elements (CATGCA) (Suzuki et al., 2007). Accordingly, when analyzing the 6-mer DNA motifs present at the proximal promoter (500 bp upstream the start codon) of the genes marked with H2AK121ub/H3K27me3 in WT and upregulated in the PRC1 mutant *atbmi1abc* (Zhou et al., 2017) (n=1030), we found an enrichment of these elements over other motifs (**Figure 1A; Supplementary Dataset 1**). In addition, VAL1/2 interact with HISTONE DEACETYLASES (HDAs)(Zeng et al., 2020; Zhou et al., 2013). Besides the B3 domain, VAL1/2 contain a Plant homeodomain like (PHD-L), a CW and an EAR domain(Suzuki and McCarty, 2008). While the PHD-L and the CW domains act as readers of H3 methylation states (Yuan et al., 2016; Hoppmann et al., 2011), the EAR domain is involved in the interaction with TPL/TRP or SAP18, which in turn recruit HDA activities (Kagale and Rozwadowski, 2011). Nevertheless, despite VAL1/2 can interact with these different repressive complexes, it is not clear whether these interactions take place simultaneously or within different contexts.

**Figure 1.**
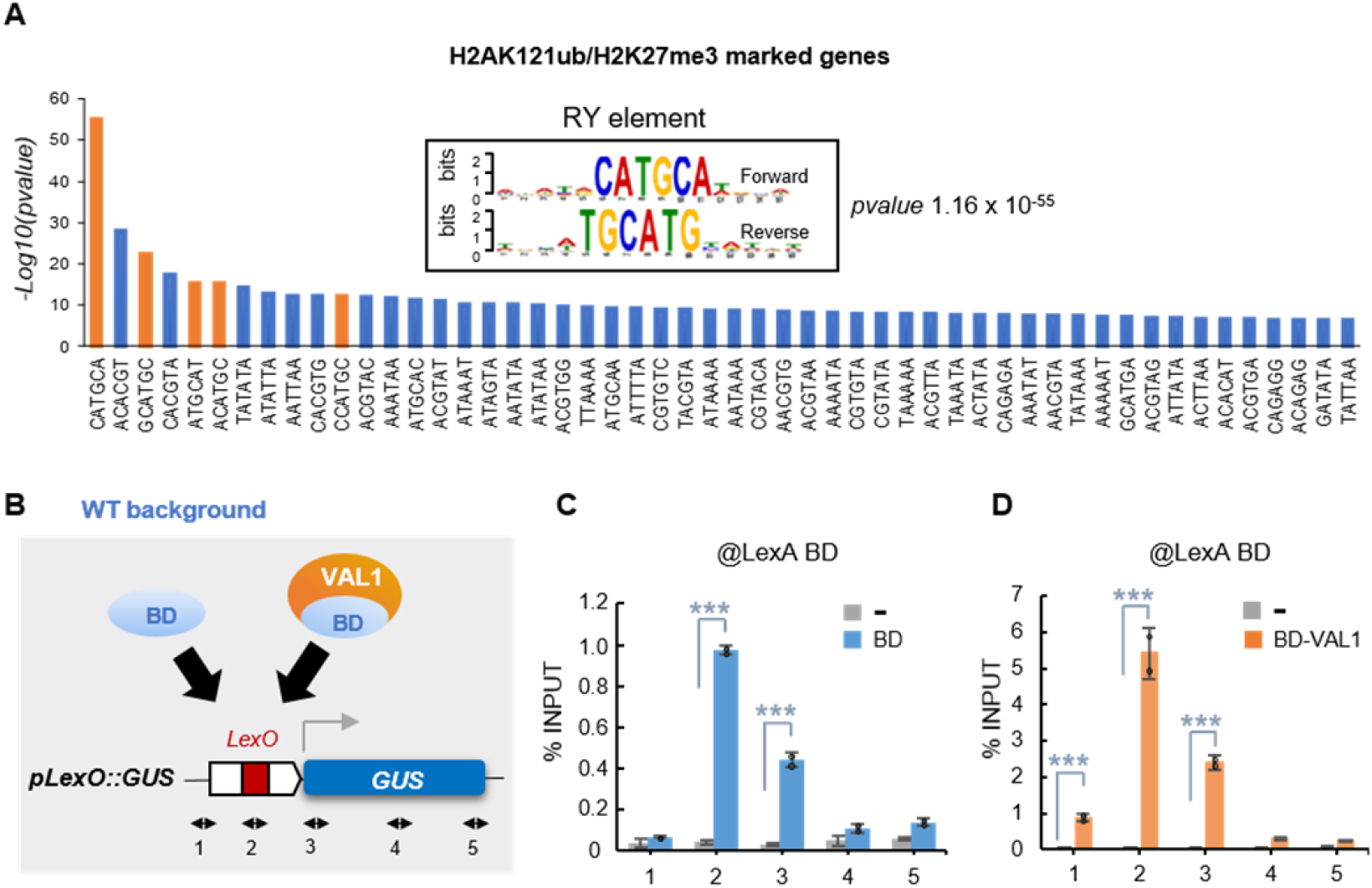
LexA BD fusion proteins *in vivo* bind to the synthetic promoter. **(A)** Bar chart showing RY element as the most significantly enriched cis-regulatory motif found at the proximal promoter of the H2AK121ub/H3K27me3 marked genes in WT that become upregulated in *atbmi1abc* mutant (n=1030 genes; see Supplementary Dataset 1). Analysis was carried out using Tair Motif finder tool (https://www.arabidopsis.org/tools/bulk/motiffinder/index.jsp). Other significantly enriched 6-mer motifs are also shown. **(B)** Schematic representation of the synthetic *GUS* reporter *locus*. The *LexO* element recognized by LexA binding domain (BD) is indicated. Numbered arrows indicate the position of the primer pairs used to examine the binding of the fusion proteins to the synthetic *locus*. **(C,D)** Bar charts showing BD (in blue) and DB-VAL1 (in orange) enrichment at *GUS* reporter *locus* determined by ChIP using anti-LexA DB antibody. WT/*pLexO::GUS* plants (in grey) were used as negative control. Results are indicated as percentage of input. Error bars represent standard deviation of n=2-3 biological replicates. Significant differences as determined by Student’s t-test are indicated (***P < 0.001).

To investigate this, we developed a system to direct VAL1 recruitment to a constitutive promoter that lacks any of the *cis* regulatory element proposed to recruit PcG activity, including RY elements. For this, we built a synthetic target promoter, consisting on a *cauliflower mosaic virus* (*CaMV35S*) promoter in which the bacterial LexA *operator* (*LexO*) was inserted. This promoter was placed upstream of the *beta-glucuronidase* (*GUS*) reporter gene, obtaining the *pLexO::GUS* construct (**Figure 1B**). In parallel, we generated a construct to express a translational fusion between LexA DNA-binding domain (BD) and VAL1, and another to express the BD alone as control (**Figure 1B**). The three constructs were independently transformed into Wild type Col-0 Arabidopsis plants (WT) and, after selecting the appropriate lines (**Supplementary Figure 1; Supplementary Figure 2**), they were crossed to obtain WT/*pLexO::GUS/BD-VAL1* and WT/*pLexO::GUS/BD* plants. To verified the functionality of the system, we confirmed the binding of the BD fusion proteins to the synthetic promoter by Chromatin Immunoprecipitation (ChIP) using anti-LexA BD antibody (**Figure 1C,D**). Next, we investigated whether H2AK121ub and H3K27me3 marks were incorporated at the reporter *locus* in the different plants. ChIP results using anti-H2AK121ub and anti-H3K27me3 antibodies showed that the incorporation of these marks was only observed in WT/*pLexO::GUS/BD-VAL1* (**Figure 2A,B**), indicating that the binding of VAL1 is able to recapitulate PRC1 and PRC2 marking. We also checked the levels of H3ac marks at the reporter *locus* in the different lines (**Figure 2C**), finding that they were considerable decreased in WT/*pLexO::GUS/BD-VAL1* compared to WT/*pLexO::GUS/BD* plants. In addition, we examined the effect of these proteins on gene expression by measuring GUS activity in the different transgenic plants (**Figure 2D**). While the binding of lexA BD did not affect the levels of GUS activity compared to control plants, the binding of BD-VAL1 led to a significant reduction of GUS activity. All together, these results indicate that VAL1 acts as a platform for the simultaneous recruitment of different histone modifying complexes involved in gene repression.

**Figure 2.**
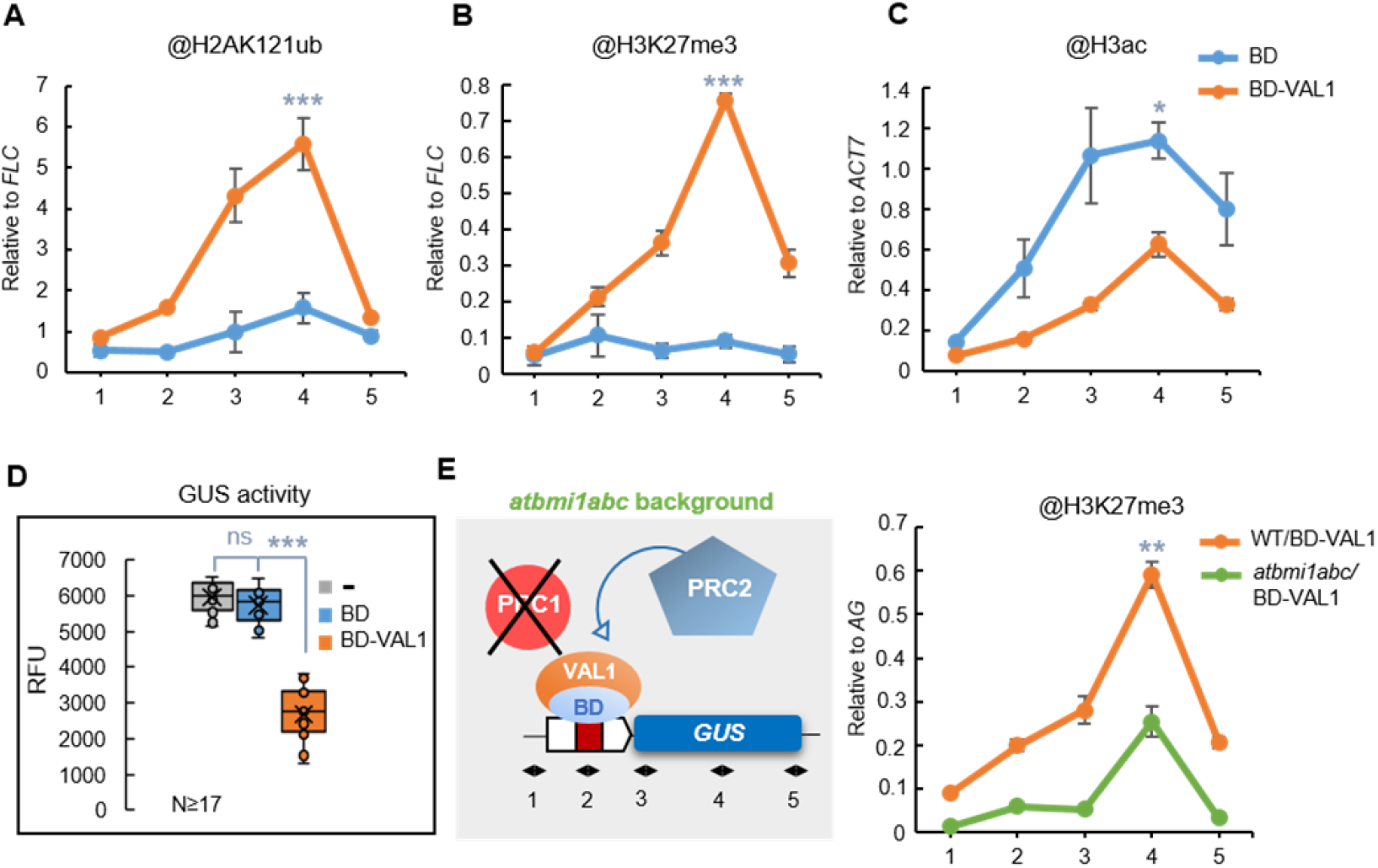
VAL1 acts as a platform for PRC1, PRC2 and HDACs recruitment. **(A,B,C)** H2AK121ub, H3K27me3 and H3ac levels at *GUS* reporter *locus* in WT/*pLexO::GUS/BD* and WT/*pLexO::GUS/BD-VAL1* plants. Numbers at *x*-axis indicate the position of amplified regions as indicated in Figure 1b. H2AK121ub and H3K27me3 levels were normalized to *FLC* and H3ac levels to *ACT7*. Error bars indicate standard deviation of n=2-4 biological replicates. Significant differences at position 4 are indicated as determined by Student’s t-test (***P < 0.001; *P < 0.05). **(D)** Box plots showing GUS activity levels in WT/*pLexO::GUS*, WT/*pLexO::GUS/BD* and WT/*pLexO::GUS/BD-VAL1* seedlings at 7 DAG. RFU indicates relative fluorescence units. Activity was tested in independent seedlings (N≥17). In each case, the median (segment inside rectangle), the mean (cross inside the rectangle), upper and lower quartiles (boxes), and minimum and maximum values (whiskers) are indicated. Significant differences as determined by Student’s t-test are indicated (***P < 0.001). “ns” indicates not significant. **(E)** Left panel, schematic representation of the experiment showed at the right panel in which the levels of H3K27me3 at the reporter *locus* were compared between WT/*pLexO::GUS/BD-VAL1* and *atbmi1abc/pLexO::GUS/BD-VAL1* plants. H3K27me3 levels were normalized to *FLC*. Error bars indicate standard deviation of n=2 biological replicates. Significant difference at position 4 is indicated as determined by Student’s t-test (**P < 0.01).

Previous reports have shown that VAL1 and AtBMI1 proteins directly interact (Yang et al., 2013; Qüesta et al., 2016) and that the levels of H3K27me3 were reduced in both *val1val2* and *atbmi1abc* mutants at seed maturation genes (Yang et al., 2013). Furthermore, genome wide analyses showed that H3K27me3 levels were reduced to some extent at most of H2AK121ub/H3K27me3 marked genes in *atbmi1abc* (Zhou et al., 2017); thus, we wondered whether VAL1 directly recruit PRC2 or if this is mediated by PRC1 interaction. To investigate this, we introduced the *pLexO::GUS* and *BD-VAL1* transgenes into *atbmi1abc* mutant (**Figure 2E**, left panel) and analyzed the levels of H3K27me3 at the reporter *locus* (**Figure 2E**, right panel). We found that despite the levels of H3K27me3 marks were considerably reduced in *atbmi1abc* mutant, they were not completely eliminated, indicating that PRC2 recruitment is mediated by both PRC1 and VAL1.

### PRC2-independent recruitment is mediated by TFs other than the VAL factors

A broad diversity of TFs belonging to different gene families are able to bind to PRE-like sequences in Arabidopsis (Xiao et al., 2017). Among the most enriched ones are the C2H2 and AP2-ERF families (Xiao et al., 2017) (**Figure 3A**). Accordingly, several evidence showed that the C2H2 factors SUP, KNU and AZF1 are able to recruit PRC2 activity (Xiao et al., 2017; Xu et al., 2018; Sun et al., 2019). Besides, the MADS-box TFs FLC and SVP have been connected to PRC2 repression (Wang et al., 2014; Richter et al., 2019) (**Figure 3A**). Therefore, we wondered whether all these TFs were able to work as recruiting platforms for PRC1, PRC2 and possibly other histone modifying complexes as VAL1 did. To test this, we generated BD-KNU, BD-FLC and BD-ERF10 fusions and analyzed their effects on our synthetic target *locus*. We found that the three fusion proteins were able to repress gene expression (**Figure 3B**) and led to the incorporation of H3K27me3 and the removal of H3ac marks (**Figure C,D**). However, we did not detect incorporation of H2AK121ub marks (**Figure 3E**). Accordingly, the *cis* regulatory motifs recognized by these TFs were enriched at the proximal promoter of the genes marked only with H3K27me3 (Zhou et al., 2017), whereas the RY elements were not enriched in this subset of genes (**Supplementary Figure 3; Supplementary Dataset 1**). These results indicate that PRC2 activity can be recruited independently of PRC1 through TFs other than VAL factors.

**Figure 3.**
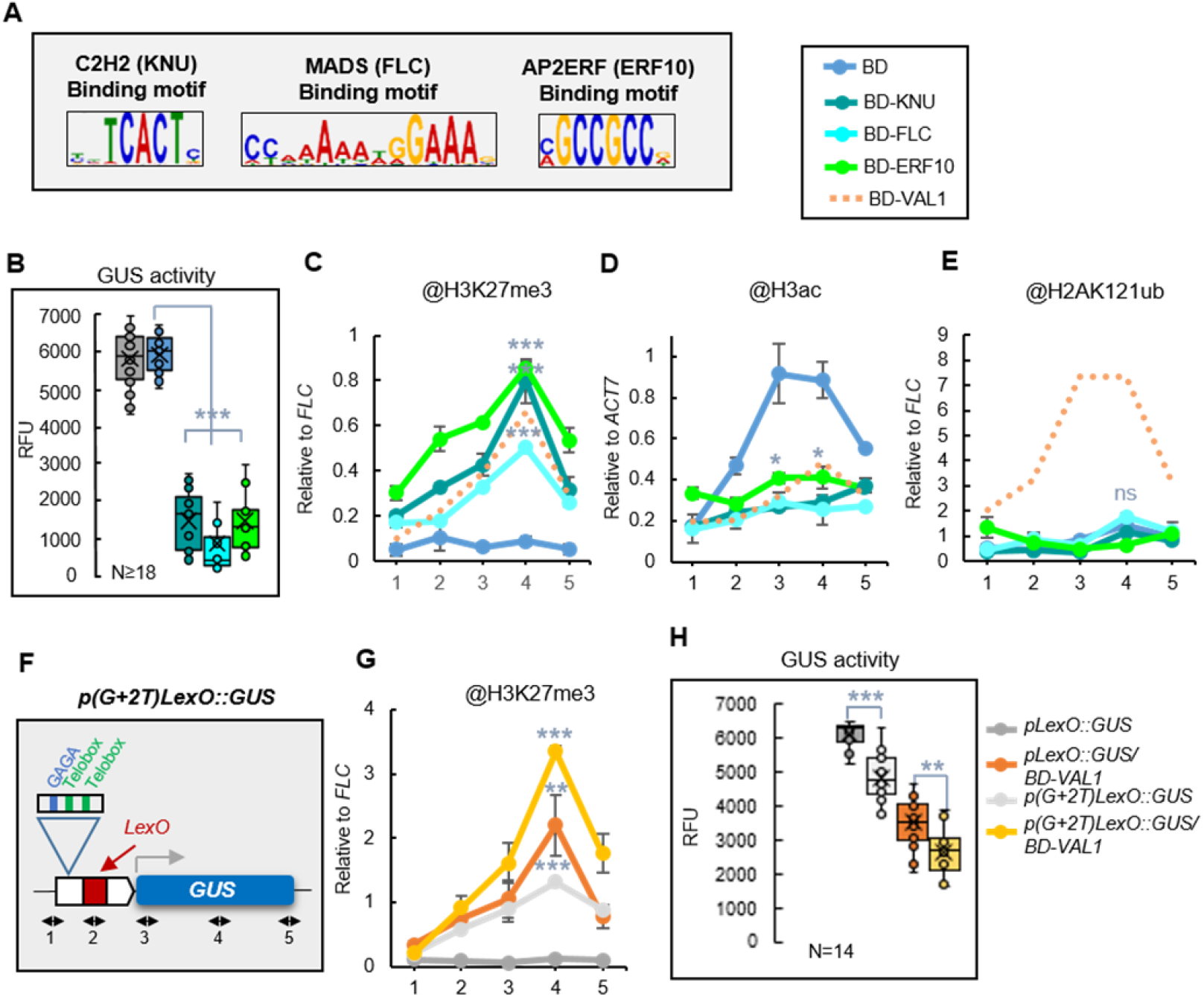
BD-KNU, BD-FLC and BD-ERF10 are able to recruit PRC2 and HDACs but not PRC1. **(A)** Sequence logos of the cis regulatory motifs recognized by C2H2, MADS and AP2ERF TFs. **(B)** Box plots showing GUS activity levels in WT/*pLexO::GUS*, WT/*pLexO::GUS/BD*, WT/*pLexO::GUS/BD-KNU*, WT/*pLexO::GUS/BD-FLC* and WT/*pLexO::GUS/BD-ERF10* seedlings at 7 DAG indicated as relative fluorescence units (RFU). Activity was tested in independent seedlings (N≥18). In each case, the median (segment inside rectangle), the mean (cross inside the rectangle), upper and lower quartiles (boxes), and minimum and maximum values (whiskers) are indicated. Significant differences as determined by Student’s t-test are indicated (***P < 0.001). **(C,D,E)** H3K27me3, H3ac and H2AK121ub levels at *GUS* reporter *locus* in the different plants. Numbers at *x*-axis indicate the position of amplified regions as indicated in Figure 1b. H3K27me3 and H2AK121ub levels were normalized to *FLC*, and H3ac levels to *ACT7*. Error bars indicate standard deviation of n=2 biological replicates. Significant differences compared to WT/*pLexO::GUS/BD* are indicated as determined by Student’s t-test (***P < 0.001; *P < 0.05). “ns” indicates not significant. One replicate of WT/*pLexO::GUS/BD-VAL1* was included as additional control. **(F)** Schematic representation of *p(G+2T)LexO::GUS* construct in which one *GAGA* and two *TELOBOX* motifs were inserted upstream of the *LexO*. **(G)** H3K27me3 levels at *GUS* reporter *locus* in WT/*pLexO::GUS* and WT/*p(G+2T)LexO::GUS* plants with and without BD-VAL1. H3K27me3 levels were normalized to *FLC*. Error bars indicate standard deviation of n=2 biological replicates. Significant differences compared to WT/*pLexO::GUS* at position 4 are indicated as determined by Student’s t-test (***P < 0.001; **P <0.01). **(H)** Box plots showing GUS activity levels in the same plants. Activity was tested in N=14 independent seedlings. Significant differences as determined by Student’s t-test are indicated (***P < 0.001; **P <0.01).

Nevertheless, since the promoters of H2AK121ub/H3K27me3 marked genes in addition to RY elements showed enrichment in other *cis* regulatory motifs (**Figure 1A; Supplementary Dataset 1**), we wondered whether different recruiting factors could collaborate in H3K27me3 marking at these genes. To test this, we inserted into the synthetic promoter a DNA fragment containing one *GAGA* and two *TELOBOX* motifs to generate the WT/*p(G+2T)LexO::GUS* line, as these motifs have been extensively related to PRC2 recruitment in Arabidopsis (Xiao et al., 2017; Zhou et al., 2018; Hecker et al., 2015b) (**Figure 3F; Supplementary Figures 1 and 4**). We first analyzed the levels of H3K27me3 at *GUS* reporter *locus* in WT/*p(G+2T)LexO::GUS* and WT/*pLexO::GUS* plants in absence of any of the BD fusion proteins. We detected some levels of H3K27me3 marks at the reporter *locus* when these motifs were present (**Figure 3G**), supporting that the TFs recognizing these motifs can mediate PRC2 recruitment. Then, we checked the levels of H3K27me3 at these reporter *loci* after the binding of BD-VAL1 (**Figure 3G**). We found higher levels of H3K27me3 in WT/*p(G+2T)LexO::GUS/BD-VAL1* than in WT/*p(G+2T)LexO::GUS*; moreover, the levels in WT/*p(G+2T)LexO::GUS/BD-VAL1* were higher than in WT/*pLexO::GUS/BD-VAL1*. Consistent with this, we found that GUS activity was lower in *p(G+2T)LexO::GUS/BD-VAL1* than in *pLexO::GUS/BD-VAL1* plants (**Figure 3H**), indicating that the levels of H3K27me3 are important to maintain gene repression. All together, these results support that different TFs can act synergistically in PRC2 recruiting.

### EAR repressome connects histone deacetylation and PRC2 marking

Since all the TFs tested, including VAL1, were able to recruit PRC2 activity, we examined if they display some common feature. Interestingly, despite the lack of sequence homology among them, these TFs contain an EAR domain. Furthermore, except for the case of TRB factors, all the TFs that have been related to PRC2 recruitment before contain an EAR domain (**Figure 4A**). The EAR domain is defined as LxLxL, DLNxP, and DLNxxP. This domain has been found in a high number of TFs of different gene families with repressive activity, constituting what has been named the EAR repressome (Kagale et al., 2010). The EAR domain mediates interaction with TPL/TPR corepressors or SAP18 protein (Kagale and Rozwadowski, 2011; Causier et al., 2012; Song and Galbraith, 2006). TPL/TPR in addition interact with the HDAs HDA6 and HDA19(Liu et al., 2014), and, importantly, with the PcG proteins EMBRYONIC FLOWER1 (EMF1) and VERNALIZATION 5 (VRN5) (Causier et al., 2012; Ke et al., 2015; Collins et al., 2019). On the other hand, SAP18 is both a component of the SIN3-HDAC (Zhang et al., 1997) and the APOPTOSIS AND SPLICING-ASSOCIATED PROTEIN (ASAP) complex (Deka and Singh, 2017). The SIN3-HDAC in Arabidopsis includes a SIN3-like protein (SNL1-6), SAP18, SAP30, one HDA activity (HDA19, HDA9, HDA7 or HDA6) and MULTICOPY SUPRESSOR OF IRA1 (MSI1)(Liu et al., 2014). Interestingly, MSI1 is also a PRC2 core component (Derkacheva et al., 2013; Mehdi et al., 2016; Ning et al., 2019). Moreover, it has been shown that SAP18 co-purifies with PRC2 core components and HDA19 (Qüesta et al., 2016). All together, these data strongly suggest a direct connection between EAR factors, TPL/TPR-HDAC or SAP18-HDAC and PRC2, which so far has not been deeply investigated. Therefore, to explore whether the EAR domain can serve as a docking point for both PRC2 and HDAC recruitment via TPL/TPR or SAP18 interaction, we compared the levels of H3K27me3 marks and H3ac at the reporter *locus* after the binding of BD-KNU or a mutated BD-KNU version in which the EAR domain was removed (BD-KNU(-EAR)) (**Figure 4B,C**). We found that the levels of H3K27me3 were significantly reduced in *pLexO::GUS/BD-KNU(-EAR)* plants compared to WT/*pLexO::GUS/BD-KNU* plants, while the opposite effect was observed for the case of H3ac. Furthermore, the levels of GUS activity in *pLexO::GUS/BD-KNU(-EAR)* were as in control plants (**Figure 4D**). To further verify these results, we checked if the LexA BD fused to an EAR domain (BD-EAR) was able to reduce H3ac, increase H3K27me3 levels and repress gene expression when recruited to the synthetic promoter. Indeed, we found that the EAR domain by itself was able to cause all these effects (**Figure 4E,F,G**); however, it was unable to recruit PRC1 (**Figure 4H**), which connects the EAR repressome to PRC2 recruitment.

**Figure 4.**
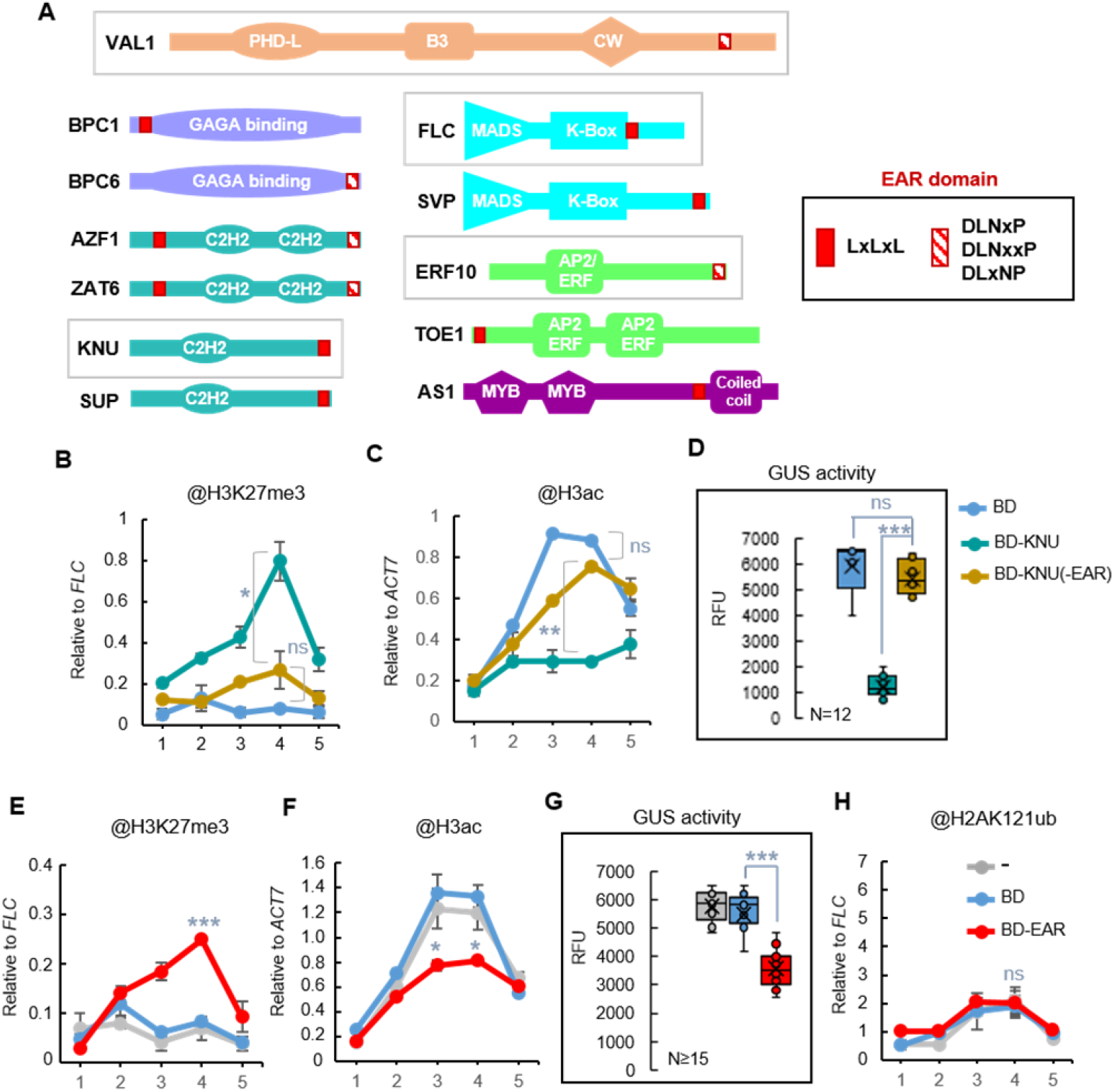
EAR domain acts as an anchoring point for PRC2 and HDACs. **(A)** Schematic representation of the domains present at TFs related to PRC2 recruitment. The TFs analyzed in this work are indicated. **(B,C)** Comparison of H3K27me3 and H3ac levels at *GUS* reporter *locus* in WT/*pLexO::GUS/BD-KNU* and WT/*pLexO::GUS/BD-KNU(-EAR)* plants. WT/*pLexO::GUS/BD* plants are included as control. Numbers at *x*-axis indicate the position of amplified regions as indicated in Figure 1b. H3K27me3 levels were normalized to *FLC* and H3ac levels to *ACT7*. Error bars indicate standard deviation of n=2 biological replicates. Significant differences at position 4 are indicated as determined by Student’s t-test (**P <0.01; *P < 0.05). **(D)** Box plots showing GUS activity levels in WT/*pLexO::GUS/BD*, WT/*pLexO::GUS/BD-KNU* and WT/*pLexO::GUS//BD-KNU(-EAR)* seedlings at 7 DAG indicated as relative fluorescence units (RFU). Activity was tested in independent seedlings (N≥12). The median (segment inside rectangle), the mean (cross inside the rectangle), upper and lower quartiles (boxes), and minimum and maximum values (whiskers) are indicated. Significant differences as determined by Student’s t-test are indicated (***P < 0.001). **(E,F)** Comparison of H3K27me3 and H3ac levels at *GUS* reporter *locus* in WT/*pLexO::GUS*, WT/*pLexO::GUS/BD* and WT/*pLexO::GUS//BD-EAR* plants. H3K27me3 levels were normalized to *FLC*, and H3ac levels to *ACT7*. Error bars indicate standard deviation of n=2 biological replicates. Significant differences between BD and BD-EAR are indicated as determined by Student’s t-test (***P < 0.001; *P < 0.05). “ns” indicates not significant. **(G)** Box plots showing GUS activity levels in the same plants indicated as relative fluorescence units (RFU). Activity was tested in independent seedlings (N≥15). Significant differences as determined by Student’s t-test are indicated (***P < 0.001). **(H)** H2AK121ub levels at the reporter *locus* in the different plants. H2AK121ub levels were normalized to *FLC*. Error bars indicate standard deviation of n=2 biological replicates.

### EMF1-TPL interaction couples H3K27me3 marking to H3 deacetylation

We also used our system to investigate the exact role of the plant-specific PcG associated factor EMF1 (Calonje et al., 2008). EMF1 has been proposed to be a PRC1 component due to its ability to *in vitro* perform similar functions to those of Drosophila Psc and to interact with AtBMI1 proteins(Bratzel et al., 2010; Calonje et al., 2008; Beh et al., 2012). However, several data indicate that EMF1 is required for H3K27me3 marking at some PcG target genes (Calonje et al., 2008; Kim et al., 2012; Li et al., 2018). Accordingly, EMF1 interacts with MSI1 (Calonje et al., 2008) and co-purifies with PRC2 components (Liang et al., 2015). In addition, EMF1 interacts with FLC and the HISTONE DEMETHYLASE JMJ14 to mediate *FT* repression (Wang et al., 2014). However, it is not clear whether EMF1 is also required for PRC1 marking. Thus, we analyzed the levels of H3K27me3 and H2AK121ub at the reporter *locus* after the binding of BD-EMF1 to the synthetic promoter (**Figure 5A**). We found that the levels of H3K27me3 were increased in WT/*pLexO::GUS/BD-EMF1* compared to WT/*pLexO::GUS/BD* plants but we did not find considerable changes in H2AK121ub levels, indicating that the recruitment of EMF1 leads to H3K27me3 marking but not to H2AK121 monoubiquitination. Since EMF1 directly interact with JMJ14 (Wang et al., 2014), we also analyzed H3K4me3 levels at the reporter *locus* (**Figure 5B**). Accordingly, we found reduced levels of these marks after BD-EMF1 binding. On the other hand, EMF1 has been shown to interact with TPL in yeast two hybrid assay (Causier et al., 2012). In support of this, we found TPL among the proteins that co-immunoprecipitate with EMF1 (Bloomer et al., 2020) (**Supplementary Dataset 2**). Furthermore, the levels of H3ac at the reporter *locus* were reduced after EMF1 binding (**Figure 5B**), confirming an EMF1-TPL-HDA interaction. We then analyzed the levels of GUS activity in WT/*pLexO::GUS/BD-EMF1* (**Figure 5C**), finding decreased levels compared to control plants; however, the levels were not as low as after the binding of the TFs, indicating that factor/s acting upstream EMF1 may be required for proper repression.

**Figure 5.**
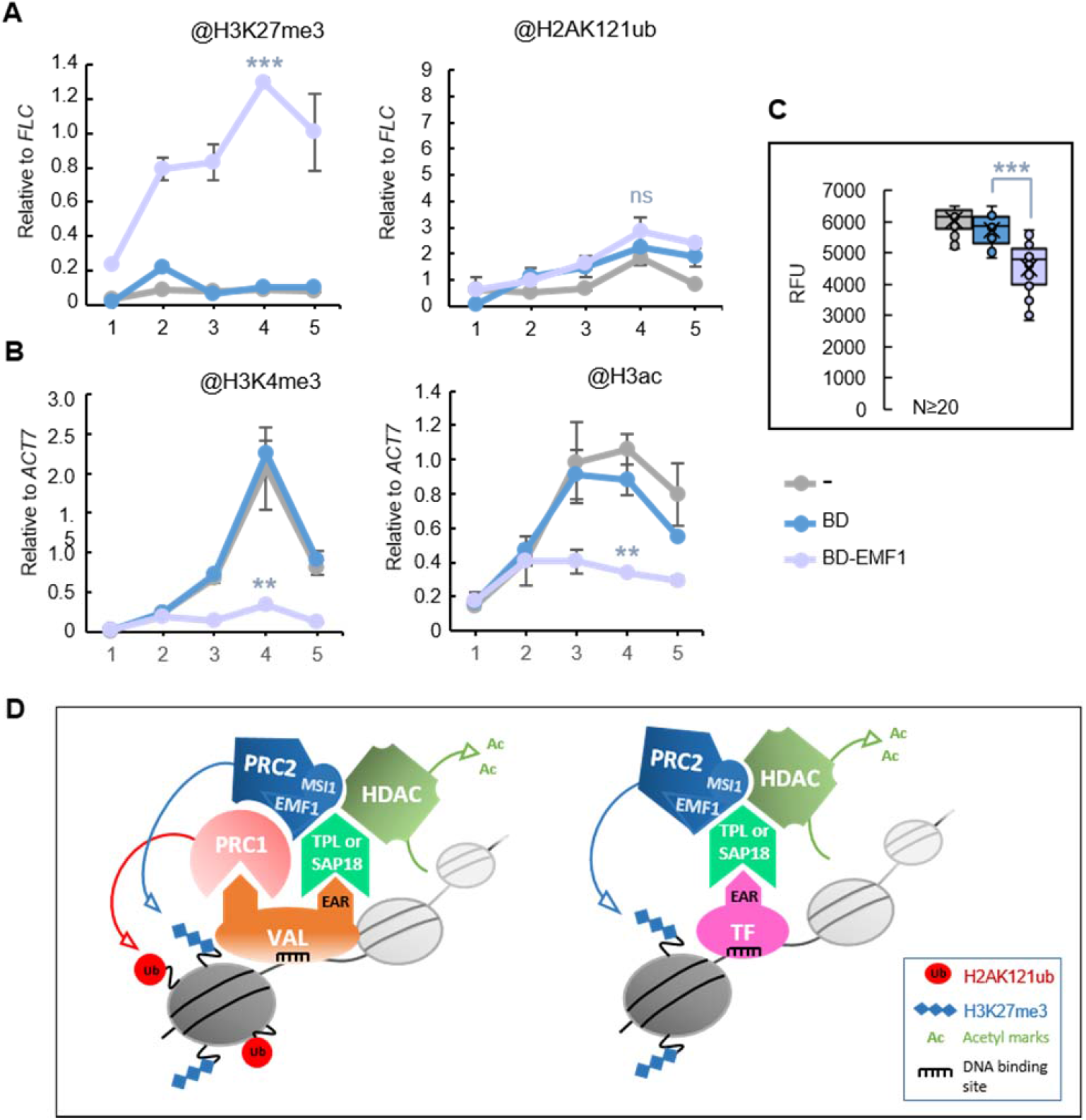
EMF1 recruitment leads to the incorporation of H3K27me3 and removal of H3K4me3 and H3ac. **(A)** H3K27me3 and H2AK121ub levels at *GUS* reporter *locus* in WT/*pLexO::GUS*, WT/*pLexO::GUS/BD* and WT/*pLexO::GUS/BD-EMF1* plants. Numbers at *x*-axis indicate the position of amplified regions as indicated in Figure 1b. H2AK121ub and H3K27me3 levels were normalized to *FLC*. Error bars indicate standard deviation of n=2 biological replicates. Significant differences compared to WT/*pLexO::GUS/BD* are indicated as determined by Student’s t-test (***P < 0.001). “ns” indicates not significant. **(B)** H3K4me3 and H3ac levels at *GUS* reporter *locus* in the different plants. The levels were normalized to *ACT7*. Error bars indicate standard deviation of n=2 biological replicates. Significant differences between BD and BD-EMF1 are indicated as determined by Student’s t-test (**P < 0.01). **(C)** Box plots showing GUS activity levels in WT/*pLexO::GUS*, WT/*pLexO::GUS/BD* and WT/*pLexO::GUS/BD-EMF1* seedlings at 7 DAG. RFU indicates relative fluorescence units. Activity was tested in independent seedlings (N≥20). In each case, the median (segment inside rectangle), the mean (cross inside the rectangle), upper and lower quartiles (boxes), and minimum and maximum values (whiskers) are indicated. Significant differences as determined by Student’s t-test are indicated (***P < 0.001). **(D)** Drawing summarizing the histone modifying complexes recruited by VAL1 or by other EAR-containing TFs to promote transcriptional repression.

## Discussion

PcG complexes ensure that each cell in an organism expresses the genes that are needed at a particular stage, time or condition. However, as PcG proteins do not have the ability to recognize DNA sequences, how PRC1 and PRC2 are recruited to the appropriate target gene is still not fully understood. Recent data support that PRC2 is recruited via interaction with different TFs; however, it is not known whether these TFs display a common feature to do so, whether the same TF can recruit PRC2 and PRC1 and how PcG-mediated differential marking is achieved. In this work, we were able to dissect how PRC1 and PRC2 recruitment take places in Arabidopsis.

We found that the binding of VAL1 TF was able to recapitulate PRC1 and PRC2 marking and to assemble HDAC activities, acting as a recruiting platform for different repressive complexes (**Figure 5D**, left panel). While PRC1 recruitment is mediated by AtBMI1 direct interaction with VAL1 (Yang et al., 2013; Qüesta et al., 2016), PRC2 recruitment involves both PRC1 and VAL1. Interestingly, our data showed that while TFs like KNU, FLC or ERF10 can mediate PRC2 and HDACs recruitment, they cannot attract PRC1 for H2A monoubiquitination (**Figure 5D**, right panel), indicating that PRC2 recruitment relies on a more general mechanism.

When comparing the protein domains present in VAL1, KNU, FLC, ERF10 and other TFs related to PRC2 recruitment before, we found that, except for the case of TRB1/2/3 factors that seem to be stable PRC2-accesory proteins as they co-purify with PRC2 (Bloomer et al., 2020), all of these TFs contain an EAR domain. The EAR domain interacts with TPL/TPR or SAP18, which in turn recruit HDA activities (Kagale and Rozwadowski, 2011; Kagale et al., 2010; Causier et al., 2012; Song and Galbraith, 2006). Interestingly, TPL and SAP18 also interact with PcG proteins. In fact, TPL co-purify with EMF1 (Bloomer et al., 2020) and SAP18 with MSI1 (Mehdi et al., 2016), suggesting that they serve as scaffolds for HDACs and PRC2 assembly. In support of this, we found that the binding of three EAR-containing TFs led to the incorporation of H3K27me3 and removal of H3ac marks at the reporter *locus*, and that this depends on the EAR domain. Moreover, the recruitment of EMF1 leads to H3K27me3 incorporation and H3ac removal. Therefore, we propose that the EAR repressome acts as anchoring point for PRC2 and HDACs recruitment (**Figure 5**).

It is unknown whether the interaction of the EAR factors with TPL/TRP or SAP18 depends on the type of EAR domain or on adjacent sequences, or whether they functionally overlap, as some of the EAR factors have been reported to interact with both (Kagale and Rozwadowski, 2011; Kagale et al., 2010; Causier et al., 2012). In any case, since TPL/TRP and SAP18 are expressed in most plant tissues (Kagale and Rozwadowski, 2011), the ability of the PcG machinery to maintain specific transcriptional states in different cell types, at different times or under different conditions, may rely on the EAR-containing recruiting factors, which expression is tightly regulated in response to internal and external signals.

We also found that different TFs can act synergistically in PRC2 recruitment at the same target gene, leading to increased levels of H3K27me3. PcG proteins in plants seems to be involved in both transient and long-term repression that persist through multiple cell divisions. Long-term repression has been reported to require spreading and maintenance of high levels of H3K27me3 marks across the target genes (Costa and Dean, 2019). Interestingly, *FLC* initial repression requires the RY elements for PcG nucleation(Costa and Dean, 2019), but its long-term repression involves other *cis* regulatory sequences located along *FLC locus* (Qüesta et al., 2020). Similarly, a PcG long-term repression in Drosophila requires sequence-specific targeting of PRC2 (Laprell et al., 2017). Thus, it might be possible that the combined action of different recruiting factors propagates and maintains appropriate H3K27me3 levels to mediate long-term repression in Arabidopsis.

Nevertheless, in Arabidopsis there is also a high number of only-H2AK121ub marked genes^16^. The promoters of these genes are highly enriched in G-box motifs (**Supplementary Figure 3; Supplementary Dataset 1**). This motif is recognized by two large families of TFs in Arabidopsis, the basic helix-loop-helix (bHLH) and Leu zipper (bZIP) families (Ezer et al., 2017), raising the possibility that TFs from these families may be involved in PRC1-independent recruitment and suggesting that PcG differential marking depends on different recruiting factors.

## Methods

### Plant material and culture conditions

*Arabidopsis thaliana* Col-0 wild type (WT), *atbmi1abc* (Yang et al., 2013) and transgenic plants harboring the different constructs were grown under long-day conditions (16 h light and 8 h dark) at 21 °C on MS agar plates containing 1.5% sucrose and 0.8% agar for 7 days. MS-agar plates were appropriately supplemented with Kanamycin (50 μg.ml-1) and/or hygromycin (10 μg.ml-1) for selection of transgenic plants.

### Synthetic system constructs and transgenic plants

To generate the synthetic target gene constructs, we used as backbone the pCAMBIA 1305.1 binary vector that contains the *GUS reported gene* under the control of the *cauliflower mosaic virus* (*CaMV35S*) promoter. We replaced the *CaMV35S* promoter by a *CaMV35S* in which the LexA DNA binding element (Lex A operator (*LexO*), amplified from pER8 vector (Zuo et al., 2000), was inserted upstream of the TATA box, resulting in the *pLexO::GUS* construct. To generate *p(G+2T)LexO::GUS* construct, a DNA fragment of 100 bp from the regulatory region of *ABSCISIC ACID INSENSITIVE 3* (*ABI3*) gene (see **Supplementary Figure 4**), which contains one *GAGA* and two *TELOBOX* motifs, was introduced into the synthetic promoter upstream of the *LexO*. These constructs were transformed into WT Col-0 plants to generate WT/*pLexO::GUS* and WT/*p(G+2T)LexO::GUS* transgenic plants. To build the BD translational fusion constructs, we inserted into the pPZP211 vector the G10-90 promoter, the LexA BD (252 bp N-terminal region of LexA protein amplified from pER8 vector (Zuo et al., 2000)), the TF cDNA and the OCTOPINE SYNTHASE (OCS) terminator. To construct the BD-EAR fusion, we used the C-terminal region of VAL1 cDNA that contains a predicted Nuclear Localization Signal (NLS) and the EAR domain (region from 2041 bp to stop codon of VAL1 cDNA; See **Supplementary Figure 5**). To ensure that the BD when expressed alone was transported to the nucleus, the sequence corresponding to VAL1 predicted NLS (region from 2041 to 2183 bp of VAL1 cDNA; See **Supplementary Figure 5**) was fused to the C-terminal region of the BD. The different BD fusion constructs were transformed into WT Col-0 plants. The expression of the protein in the different transgenic lines was verified by Western blot using anti-LexA BD antibody. One line from each BD fusion was crossed to the same WT/*pLexO::GUS* or WT/*p(G+2T)LexO::GUS* line. Primers used are listed in **Supplementary Dataset 3**.

### Western blot assay

Total protein extract from the different plants was separated on 10% SDS-PAGE gel and transferred to a PVDF membrane (Immobilon-P Transfer membrane, Millipore) by semi-dry blotting in 25 mM Tris–HCl, 192 mM glycine, and 10% methanol. To detect the fusion proteins, anti-LexA BD antibody (Millipore 06-719; 1:2000) was used as primary antibody and Horseradish peroxidase-conjugated goat anti-rabbit (Sigma-Aldrich, A0545; 1:10,000) as secondary. Chemiluminescence detection was performed with ECL Prime Western Blotting Detection Reagent (GE Healthcare Life Sciences) following the manufacturer’s instructions.

### Fluorometric assay of beta-glucuronidase (GUS) activity

The activity of beta-glucuronidase (GUS) was determined on whole seedlings using 4-methylumbelliferyl ß-D-glucuronide (4-MUG) as a substrate (Halder and Kombrink, 2015). One-Single 7-day-old seedlings were placed in 96-well microplates and incubated with 150 μL lysis buffer (50 mM sodium phosphate, pH 7.0, 10 mM EDTA, 0.1% Triton X-100) containing 1mM 4-MUG at 37°C for 90 min. At the end of the incubation period, 50 μL of 1M Na2CO3 (stop solution) was added to each well and 4-MU fluorescence was directly determined in a microplate reader (excitation/emission wavelength of 365/455 nm). Activity is expressed as relative fluorescence units (RFU).

### Chromatin immunoprecipitation (ChIP) and ChIP-qPCR

ChIP experiments were performed on one gr of 7-day-old whole seedlings fixed in 1% Formaldehyde. Chromatin was extracted from fixed tissue and fragmented using a Bioruptor^®^ Pico (Diagenode) in fragments of 200-500 bp. The sheared chromatin was immunoprecipitated overnight using the following antibodies and dilutions: anti-LexA BD (Millipore 06-719, 1:300) Anti-H3K27me3 (Millipore, 07-449, 1:300), anti-H2AK121ub (Cell Signaling, 8240S; 1:100) Anti-Histone H3 (acetyl K9 + K14 + K18 + K23 + K27) (Abcam ab47915, 1:300), or anti-H3K4me3 (Diagenode, C15410003-50; 1:300). Immunocomplexes were captures using Protein-A Sepharose beads CL-4B (GE Healthcare). After washing the Protein-A beads, chromatin was eluted and the cross-linking was reversed overnight at 65°C. The DNA from the immunoprecipitated chromatin was treated with RNase and proteinase K and purified by phenol-chloroform extraction followed by ethanol precipitation. For ChIP-qPCR, amplification was performed using SensiFAST™ SYBR^®^ & Fluorescein Kit (Bioline) and iQ5 Biorad system. Results are given as percentage of input or as relative level to *FLOWERING LOCUS C* (*FLC*), *ACTIN 7* (*ACT7*) or *AGAMOUS* (*AG*), depending on the histone mark and the genotype. qPCR data are shown as the means of two to four biological replicates as indicated. Primers used for ChIP-qPCR are listed in **Supplementary Dataset 3**.

## Author Contributions and Acknowledgements

FB and WM performed the experiments with the help of IH. MC designed the study, interpreted the results and wrote the manuscript.

This work is supported by BIO2016-76457-P, PID2019-106664GB-I00 Grants from Spanish Ministry of Science and innovation. IH was supported by a Spanish National Research Council (CSIC) training scholarship (JAEINT2018-EX-0821).

